# Flow over seal whiskers: importance of geometric features for force and frequency response

**DOI:** 10.1101/2020.04.13.039750

**Authors:** Kathleen Lyons, Christin T. Murphy, Jennifer A. Franck

**Affiliations:** Department of Engineering Physics, University of Wisconsin–Madison, Madison WI; Naval Undersea Warfare Center, Newport RI

## Abstract

The complex undulated geometry of seal whiskers has been shown to substantially modify the turbulent structures directly downstream, resulting in a reduction of hydrodynamic forces as well as modified vortex-induced-vibration response when compared with smooth whiskers. Although the unique hydrodynamic response has been well documented, an understanding of the fluid flow effects from each geometric feature remains incomplete. In this computational investigation, nondimensional geometric parameters of the seal whisker morphology are defined in terms of their hydrodynamic relevance, such that wavelength, aspect ratio, undulation amplitudes, symmetry and undulation off-set can be varied independently of one another. A two-factor fractional factorial design of experiments procedure is used to create 16 unique geometries, each of which dramatically amplifies or attenuates the geometric parameters compared with the baseline model. The flow over each unique topography is computed with a large-eddy simulation at a Reynolds number of 500 with respect to the mean whisker thickness and the effects on force and frequency are recorded. The results determine the specific fluid flow impact of each geometric feature which will inform both biologists and engineers who seek to understand the impact of whisker morphology or lay out a framework for biomimetic design of undulated structures.

## Introduction

Harbor seals, among other seal species, demonstrate incredible hydrodynamic tracking abilities [1]. The advanced tracking ability of the seal is attributed in part to the unique morphology of its whiskers. The undulated morphology is shown to significantly reduce hydrodynamic forces as well as modify the vortex-induced-vibration (VIV) response when compared to smooth whiskers [2]. As shown in Fig 1, the regular shedding pattern of low pressure, alternating sign vortices on a smooth elliptical cylinder is replaced with a complex structure of intertwined vorticity when the seal whisker undulations are added. These flow structures create a higher pressure region immediately downstream, reducing drag force and lift force oscillations. Understanding the complex fluid dynamics and frequency response of whisker undulations will aid in understanding the impact of seals’ specialized whisker morphology, and help provide context for differences observed between species. Furthermore, biomimetic inspiration from the whisker undulations could impact a multitude of engineering designs that seek to minimize VIV, reduce drag, or disrupt the frequency response across a large variety of structures such as mooring ropes and wind turbine bases.

**Fig 1.**
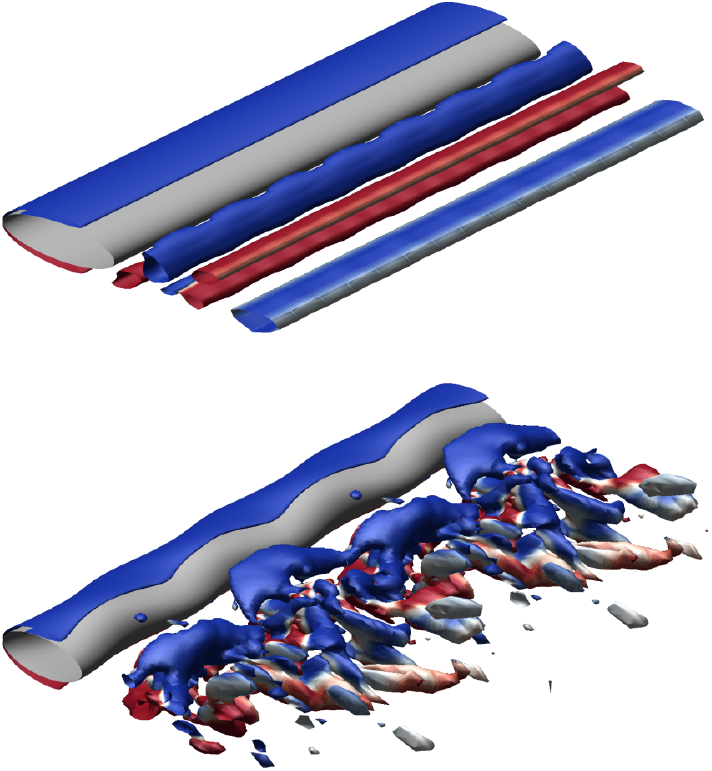
Visualization of hydrodynamic effects of seal whisker geometry. Isosurfaces of nondimensional *Q*-criterion value 0.4, colored by positive (red) and negative (blue) *z*-vorticity. Flow over a smooth ellipse (top) compared with the flow over the seal whisker model (bottom) reveal distinctly different flow structures and hydrodynamic responses as a result of the undulations.

The tracking performance of seal whiskers is well documented through a series of hydrodynamic-trail following exercises with trained blind-folded seals [1, 3]. Through these experiments, it is shown that harbor seals can detect flow disturbances with their whiskers, even those generated up to 40 meters away. Additionally, seals are shown to distinguish between disturbances left by objects of different sizes and shapes [4], demonstrating that they can use their whiskers to discriminate among specific flow signatures, and likely among various prey types. To complement the hydrodynamic tracking experiments, systematic exploration of the whisker system’s tactile sensitivity in a live seal reveals the ability to detect perturbations on the order of 1 mm/s [5]. In controlled laboratory flume experiments, excised seal whiskers exhibit a broad range of frequency response in water flow with amplitudes noticeably influenced by the angle of attack, or orientation of the whisker, with respect to the freestream flow [6, 7].

To better understand how the whisker geometry enables this response sensitivity, the harbor seal whisker is modelled as shown in Fig 2. Described by Hanke et al. in 2010 [2], this model geometry is defined by two ellipses with centers a distance *M* apart, where one is more oblique and the other more circular in shape. The ellipses are defined by their major radii, *a* and *k*, and minor radii, *b* and *l*. Both ellipses are tilted with respect to the *x*-axis at incident angles *α* and *β* respectively, which results in a spanwise offset between the leading and trailing edge undulations. Collectively, these features create a complex three-dimensional and spanwise-periodic whisker model. The fluid flow is nominally positioned in the positive *x*-direction, along the more streamlined coordinate axis of the whisker. Results from a combination of experimental particle-image velocimetry (PIV) and computational fluid dynamics (CFD) experiments show that the spanwise undulations of the seal whisker model disrupt the strong von Kármán vortex street that is characteristic of smooth cylinders, thus modifying the downstream vortical structures and resulting forces [2]. The study by Hanke et al. among others, demonstrates effective suppression of VIV in the seal whisker model.

**Fig 2.**
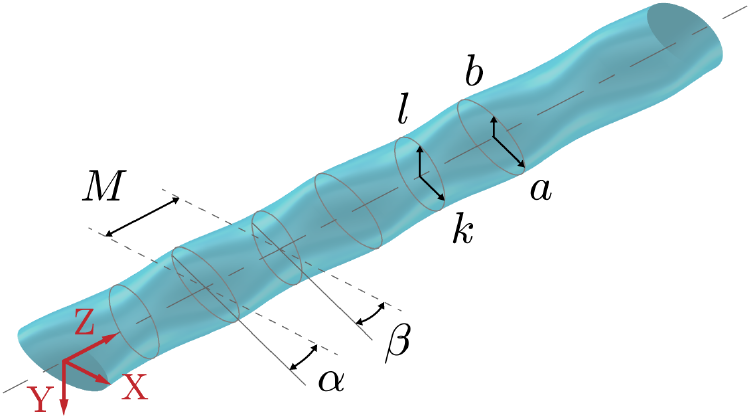
Geometry of baseline model. Geometric parameters of the baseline model as originally defined by Hanke et al. [2] create two coordinating sets of spanwise undulations along the *z*-axis. The seal whisker model is created from two different sized ellipses a distance *M* apart, major radii *a* and *k*, minor radii *b* and *l*, inclined at incident angles *α* and *β* with respect to the *x*-axis. Nominal values for harbor seal whiskers are *M* = 0.91 mm, *a* = 0.595 mm, *b* = 0.240 mm, *k* = 0.475 mm, *l* = 0.290 mm, *α* = 15.27 degrees, and *β* = 17.60 degrees [2].

The whisker model in Fig 2 along the with nominal values published by Hanke et al. continue to be the subject of both computational and experimental investigation especially surrounding biologically relevant flow regimes. At *Re* = 1800, a direct comparison to an elliptical cylinder and a simple wavy cylinder is performed experimentally, where dynamic mode decomposition of the flow field indicates strong disruption of the energy redistribution process immediately downstream of the whisker [8], which varies with respect to angle of attack [9, 10]. The drag reduction properties are further confirmed computationally via a lattice Boltzmann method, in which a hydrodynamic disturbance is created via a paddle positioned 12.1 diameters upstream to assess the whisker’s response to stimulation [11].

The whisker model described above is constructed from average measurements of digital photography from 13 adult harbor seal whiskers, which exhibit considerable variation in some geometric parameters, specifically the angles of incidence *α* and *β* [12]. A follow-up investigation by Rinehart et al. [13] performs independent whisker characterization using 27 whiskers from harbor and elephant seals. Among the findings are smaller mean angles of incidence, *α* and *β*, of 0.303 and 1.079 degrees respectively, in contrast with the larger angles presented by Hanke et al. of *α* = 15.27 degrees and *β* = 17.60 degrees. However, both investigations note the large statistical variability in the angles, which directly affect the amplitude of the undulations on the seal whisker. Research on the fluid flow over whisker models has varied in the representation of the angles of incidence with some research claiming they are relatively unimportant features [14] and others setting the angles of incidence to zero [12, 15]. In addition to the aforementioned models primarily built from harbor seals and elephant seals, morphological analyses report geometric variation both within and between species [16, 17]. The statistical variation found in nature highlights the relevance of exploring the hydrodynamic effects of geometric variation in the seal whisker morphology.

The model parameters proposed by Hanke et al. are most prevalent throughout the literature and only a handful of geometry variations are explored computationally. Although the flow over the seal whisker system has been extensively studied, a systematic investigation where all geometric parameters are independently modified has never been completed. Witte et al. explore two variations that indicate a larger wavelength has a slightly decreased root-mean-square (RMS) lift force but the same mean drag force [12]. Hans et al. explore four different geometric models and demonstrate that both coordinate directions of undulation amplitudes are necessary for maximum drag-reduction benefits, however, the angle of incidence has only a weak influence on drag [14]. In a recent work, Liu et al. [15] investigate a geometric model based on the parameter values presented in the work of Rinehart et al. [13]. Results indicate that the undulations in opposing directions must alternate with one another along the span of the seal whisker for optimal hydrodynamic force reduction. By modifying the amplitude and wavelength, they show that the whisker morphology suppresses lift oscillation over a wide range of parameters. However, without systematic and substantial variations in undulation morphology, it remains unclear which geometric features or combinations of features are most important, and how specific features correlate with force reduction and frequency response.

Closely related to the seal whisker geometry is that of a wavy cylinder, which has been more thoroughly investigated in literature [18–21]. The wavy cylinder geometry is defined by a sinusoidally varying diameter, *D*(*z*), along its spanwise direction prescribed by

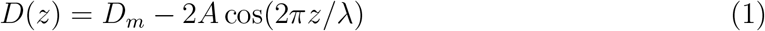

where *D*_*m*_ is the average diameter, *A* is the amplitude of the variation, *λ* is the wavelength, and *z* is the distance along the spanwise length. In wind tunnel experiments at *Re* = 30, 000 completed by Lam et al., a wavy cylinder is shown to produce a three-dimensional separation line in the boundary layer [19]. Importantly, Lam et al. also include amplitude variation in their study, noting that larger *obliqueness*, defined as *A*^2^/(*λD*_*min*_), results in higher levels of boundary separation and pressure effects due to three-dimensionality [19]. Through the use of PIV in a circulating water channel at *Re* = 3000, Zhang et al. show that the geometry of the wavy cylinder can reduce drag and lift. They hypothesize that the counter-rotating secondary vortices that are formed on either side of maximum diameter locations suppress the formation of larger-scale spanwise vorticity [20]. In a numerical investigation, Lam and Lin assess the effects of a range of amplitudes and wavelengths on the force reduction properties of wavy cylinders at *Re* = 100 [21]. The simulations include a range of wavelength ratios, *λ/D*_*m*_, from 1 to 10 and a range of amplitude ratios, *A/D*_*m*_, from 0.050 to 0.250. A comparison of the average drag coefficients and the RMS lift coefficients for the models show nonlinear trends, containing both local minima and maxima for the force coefficients within the range of *λ/D*_*m*_. Lam and Lin classify three basic flow patterns based on wavelength. At low wavelengths, flow structures appear similar to those of smooth cylinders; at moderate wavelength three-dimensional distortion and vortex formation length are increased; and at large wavelength, the free shear layer tends to not roll up at all.

One can gain insight into the flow physics over the whisker topography from previous wavy cylinder research. Although both have spanwise undulations, the whisker geometry has additional complexity. Whereas the wavy cylinder geometry is described by two variables, amplitude and wavelength, the whisker geometry can only be fully described by no less than seven parameters as shown in Fig 2. These additional variables describe the size, orientation, and relative location of the two sets of undulations that are present in the whisker geometry in contrast to the single undulation that governs the wavy cylinder geometry. The amplitude and wavelength of the wavy cylinder geometry have been well studied allowing for creation of a response surface with respect to both parameters. Given the large number of parameters required to define the whisker geometry, a similar full response surface with respect to all parameters would be impractical. Nevertheless, the 16 whisker models evaluated in the current investigation comprise the largest geometry modification study to be completed to date to the authors’ knowledge.

The investigation presented here assesses the relative importance of each geometric parameter through systematic modifications. By intensifying and modifying the geometric features, their effects and interactions are more readily observed, and a better understanding of their contribution to drag reduction, RMS lift reduction, and frequency response modification can be achieved. First, new nondimensional, independent geometric parameters are defined by hydrodynamic relevance to enable systematic variation. Each of these parameters is varied dramatically from the baseline model, beyond the range of what is biologically found in seals.

A 16 simulation test matrix of various seal whisker topographies, each with geometry modifications derived from the model proposed by Hanke et al. [2], is created using a two-factor fractional factorial design of experiments. The parametric study of various morphological features is performed computationally with a large-eddy simulation (LES) at *Re* = 640 based on hydraulic diameter and freestream velocity, where the choice of Reynolds number is motivated by biological relevance. Reasonable foraging speed for harbor seals ranges from approximately 0.5 to 2 m/s [22], corresponding to Reynolds numbers between roughly 300 and 1400. The scope of this study is focused on investigating the importance of geometric parameters, therefore, each computation simulates an infinite length whisker model. By neglecting whisker taper and tip effects, the topography modifications can be isolated. The resulting time-averaged forces and frequency spectra are recorded, as well as velocity contours and streamlines at various spanwise locations along the whisker model.

The purpose of this investigation is to assess parameter significance rather than to develop a highly resolved response surface with respect to all features. A reduced parameter space provides an opportunity for more focused high resolution analysis in the future. Furthermore, isolation of individual feature effects provides information for the biological community to put morphology differences between species [16, 17] into context. The following section will describe the modifications of the geometric features and detail the computational methods. The results and discussion will highlight the most important geometric features and two-feature combinations affecting the forces and frequency and then discuss the resulting flow structures and downstream flow characteristics.

## Methods

### Geometric Parameters

The seal whisker model geometry in Fig 2 is defined by seven geometric parameters as previously described in the Introduction. Although defining the model based on the relationship of two ellipses (using the parameters *M* . *a*, *b*, *k*, *l*, *α* and *β*) is convenient for constructing the model it is difficult to make systematic variations and isolate the effects of the major/minor radius lengths and angles of incidence as both of these parameters influence the undulation amplitudes. Thus, using the same model, the geometric parameters are redefined in Fig 3. A summary of the new geometric descriptions and their nondimensionalization is given in Table 1.

**Fig 3.**
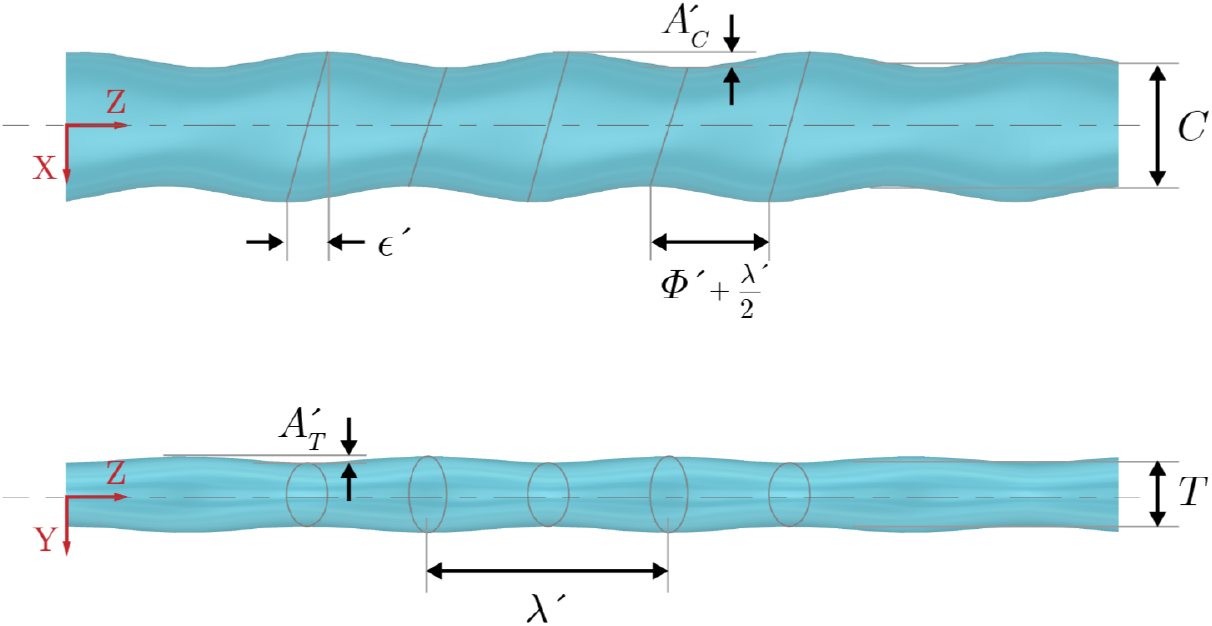
Schematic of baseline whisker model. Schematic shows the hydrodynamic-based geometric parameters that define the undulation features as described in Table 1. Top view (top panel) and front view (bottom panel), where flow is along the positive *x*-axis.

**Table 1.**
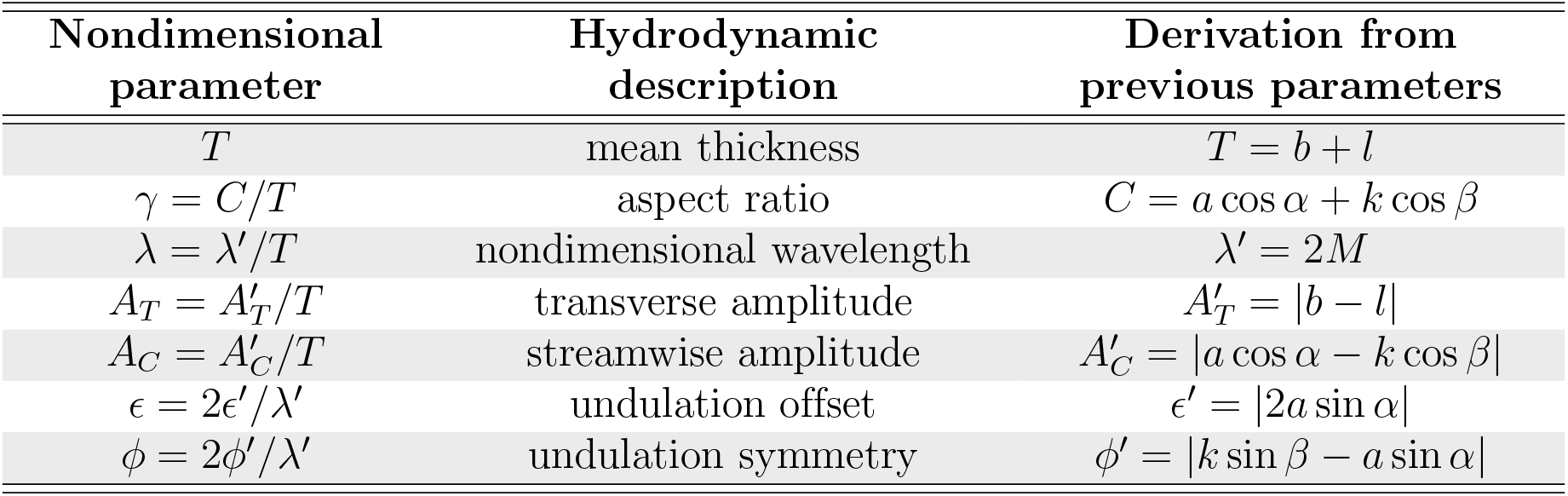
Description of geometric parameters. The description of the undulated geometric features shown in Fig 3 and the correspondence between those previously defined by Hanke et al. [2] (shown in Fig 2).

Defining freestream flow in the positive *x*-axis and following aerodynamic convention, the two major length parameters in Fig 3 are the mean thickness, *T*, and the mean chord length, *C*. The aspect ratio, or slenderness, of the geometry, is defined as *γ* = *C/T*. The amplitudes of undulations in thickness and chord length are described by *A*_*T*_ and *A*_*C*_ respectively. The periodic occurrence of both the chord and thickness undulations is defined by a single wavelength, *λ*. The wavelength and amplitudes are nondimensionalized by the mean thickness, *T*.

The undulation offset, *ϵ*, controls the alignment of undulation peaks along the leading and trailing edges, as measured in the *x-z* plane of Fig 3. When *ϵ* = 0, the peak amplitudes from leading to trailing edge are aligned with the *x*-axis. The undulation symmetry, *ϕ*, defines the steepness, or bias, of the undulation. The dimensional value of *ϕ*′ is best represented in the *x-z* plane in Fig 3 where it depicts the distance by which the trough of the undulation has been shifted, or biased. When *ϕ* = 0, the undulations along the leading and trailing edges are exactly sinusoidal. The undulation offset and symmetry parameters are nondimensionalized by the half-wavelength, *λ*′/2.

### Simulation Matrix of Geometric Parameter Combinations

Obtaining a full response surface with respect to all parameters would require an impractical number of simulation cases. Even a coarse response curve, simply five data points, with respect to each of the six nondimensionalized parameters would result in a parametric study with a matrix of 5^6^ = 15, 625 parameter combinations. Rather than obtain a full response surface *a priori*, it is evident that reduction of the parameter space is necessary to grasp a fundamental understanding of the system and support focused analysis in the future. Using a two-parameter fractional factorial design, similar to the design structure used by Jung et al. [23], a matrix of simulations is constructed to independently test the influence of each geometric parameter. The parameter values outlined in Hanke et al. [2] represent the baseline dimensions for the analysis, hereafter referred to as the *baseline model*. These values are used throughout the literature as representative values for the seal whisker geometry [8–11, 14, 24, 25]. In order to understand the effects of each nondimensional parameter, values above and below the baseline value are outlined in Table 2. A total of 16 simulations are performed with a specific combination of the low (−1) and high (1) values. The resulting test matrix, informed by the fractional factorial design of experiments, is designed as a screening test of specific combinations that systematically reduces the number of cases required to determine the most significant variables [26, 27]. Geometric parameter significance is evaluated across the tested range and is independent of any nonlinearity within the given range. Using this combination of simulations, the geometric parameters and two-parameter interactions that most strongly influenced the drag, the root-mean-square lift, and the dominant frequency response are identified and ranked by importance. This methodology allows for identification of the most influential geometric parameters rather than producing a full response surface with respect to each one. After the most significant parameters have been identified, the full response curves can be mapped in future investigations.

**Table 2.** Low and high geometric parameter values. The low and high values used to vary each of the geometric parameters compared to the baseline values commonly used in literature. The full matrix of simulations is described in detail in Table 5.

| | $\lambda$ | $\gamma$ | $A_T$ | $A_C$ | $\epsilon$ | $\phi$ |
| --- | --- | --- | --- | --- | --- | --- |
| Low (-1) | 1.8 | 1.0 | 0.05 | 0.05 | 0.20 | 0 |
| Baseline (0) | 3.4 | 1.9 | 0.09 | 0.23 | 0.34 | 0.015 |
| High (1) | 5.0 | 3.0 | 0.30 | 0.30 | 0.60 | 0.300 |

To amplify the effect on the response variables, geometric high and low values are chosen to span a wide range of deviation from the baseline model. Values of *λ* are influenced by previous research on wavy cylinders that show an increase in forces when *λ/D*_*m*_ < 2, where *D*_*m*_ is the mean diameter [21]. Therefore, the low value was chosen to be *λ* = 1.8 and the high value at an equal Δ*λ* above the baseline at *λ* = 5. The aspect ratio ranges from a circular cross section (*γ* = 1) to an elliptical cross-section of *γ* = 3, again maintaining approximately the same Δ*γ* in each direction from the baseline value. The amplitudes, *A*_*T*_ and *A*_*C*_, have low values of 0.05, or 5% of the thickness, such that they are small but nonzero. The high value of both amplitudes is 0.3, or 30% of the thickness, which represent distinct undulations. The low value of *ϕ* is zero, representing perfectly symmetrical undulations, whereas the high value is 0.30, or a 30% bias in the undulation symmetry. The low value of the undulation offset is *ϵ* = 0.2 to allow for a *ϕ* value of zero to be reached for all configurations. The high *ϵ* value is 0.6, representing a very noticeable phase difference between the undulations on the leading and trailing edges.

The effects of each geometric parameter are assessed with three response variables. The drag and lift coefficients are computed as,

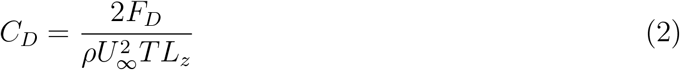

and

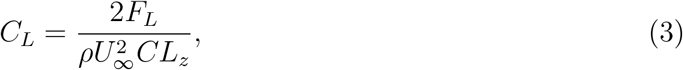

where *L*_*z*_ is the span length in the *z*-direction. Drag is normalized by the averaged frontal area, *TL*_*z*_, and lift is nondimensionalized by the planform area, *CL*_*z*_. Due to the lack of camber in all models the mean *C*_*L*_ is nominally zero, and thus the mean drag coefficient, 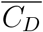, and the root-mean-square (RMS) value of the lift oscillations, *C*_*L,RMS*_ are reported as response variables. The frequency response is obtained from the oscillating lift coefficient, and is nondimensionalized as

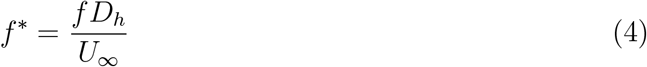

where *D*_*h*_ is the average hydraulic diameter of the model calculated in the same manner as Witte et al. [12]. The frequency response variable is the Strouhal number (*St*), which is defined in this context as the nondimensional frequency with the strongest response peak.

The average effect of each parameter on the response variable is calculated by computing the difference between the average of the response to the parameter at the high level and the average of the response to the parameter at the low level [26, 27]. The average effect of a parameter *A* on the response variable 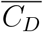 is

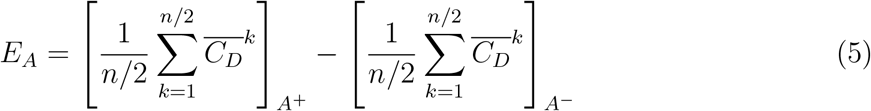

where *A*^−^ represents all cases with a low value of *A* and *A*^+^ represents all cases with a high value. For a matrix of *n* = 16 simulations, the response variable output for the *k*^*th*^ simulation is given by 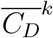. The effect of two-way interactions of geometric parameters, defined as *A*\**B*, is similarly

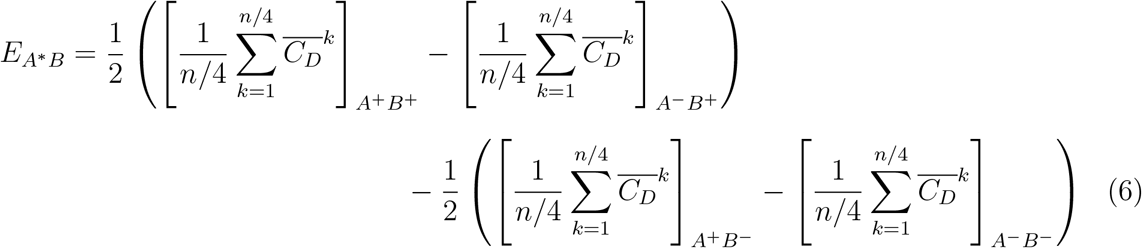

To illustrate flow structures, isosurfaces of nondimensional *Q*-criterion are plotted. *Q* is defined as the second invariant of the velocity gradient tensor, *∂u*_*i*_/*∂x*_*j*_, calculated from the vorticity and rate-of-strain tensors [28] as

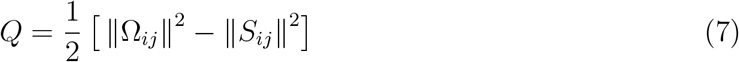

where Ω_*ij*_ and *S*_*ij*_ are defined as

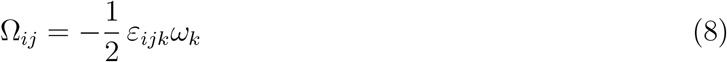

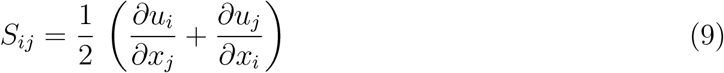

and *ω*_*k*_ represents values of the vorticity vector components. Using this definition, *Q* is positive when rotation dominates and negative when strain dominates. The criterion *Q* > 0 is frequently used to identify the location of vortex structures especially when paired with overlapping regions of low pressure, *p* < *p*_∞_, such as those that occur in the whisker’s wake.

In this investigation, *Q* is nondimensionalized by mean thickness, *T*, and freestream velocity, *U*_∞_. The isosurface value plotted is chosen to best illustrate the vortex structures and enable comparison between various models. Elliptical aspect ratio models are shown with isosurfaces of *Q*/(*U*_∞_/*T*)^2^ = 0.8 and circular aspect ratio models are shown with isosurfaces of *Q*/(*U*_∞_/*T*)^2^ = 1.6.

### Computational Details

An incompressible large-eddy simulation (LES) is used to perform the simulations by solving the filtered Navier-Stokes equations,

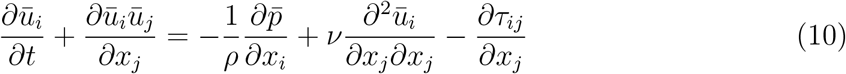

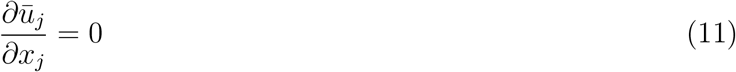

where overbar represents a low-pass spatially filtered quantity, *u*_*i*_ are the three components of velocity, *p* is pressure, *ν* is kinematic viscosity, and *ρ* is density. The sub-grid scale stress term, *∂τ*_*ij*_/*∂x*_*j*_, is calculated using a constant Smagorinsky model [29] where 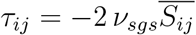. The sub-grid scale viscosity, *ν*_*sgs*_, is modelled as 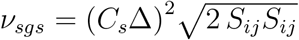 with a Smagorinsky constant of *C*_*s*_ = 0:134 where Δ is the mesh grid size.

The governing equations are solved using OpenFOAM libraries [30] that utilize a second-order accurate finite-volume scheme using Gaussian integration and linear interpolation from cell centers to cell faces. In order to enhance numerical stability and increase the time-step, a mixed semi-implicit method for pressure linked equations (SIMPLE) and pressure-implicit split-operator (PISO) algorithm is implemented with two outer correction loops. The time-stepping routine is a second-order accurate backwards method, and a stabilized preconditioned conjugate gradient method solves the matrix equations.

In order to sweep through a large matrix of 16 different three-dimensional geometries, much care is taken to reduce the computational cost yet still maintain an accurate description of the salient features of the geometry, particularly in terms of properly resolving the response variables 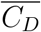, *C*_*L,RMS*_, and *St*. The final mesh for the baseline model is depicted in Fig 4, and contains 160 points in the circumferential direction, 68 points in the radial direction, and 68 points in the spanwise direction. The outer domain is circular, and is 75*T* away from the centered body. Two wavelengths are simulated in the spanwise direction with a periodic boundary condition imposed on each end. Each whisker model is centered at the origin oriented perpendicular to the flow at zero angle of attack. The freestream flow is in the positive *x*-direction and for *x* < 0, the outer boundary is subject to inlet conditions with constant velocity and zero pressure gradient. Outlet boundary conditions of zero velocity gradient and constant pressure are applied for *x* > 0. No-slip boundary conditions are applied at the surface of the whisker model. The values for the response variables 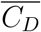, *C*_*L,RMS*_, and *St* are calculated over 700 convective time units. The final mesh selection for the baseline model produces results that compare favorably with a high resolution direct numerical simulation (DNS).

**Fig 4.**
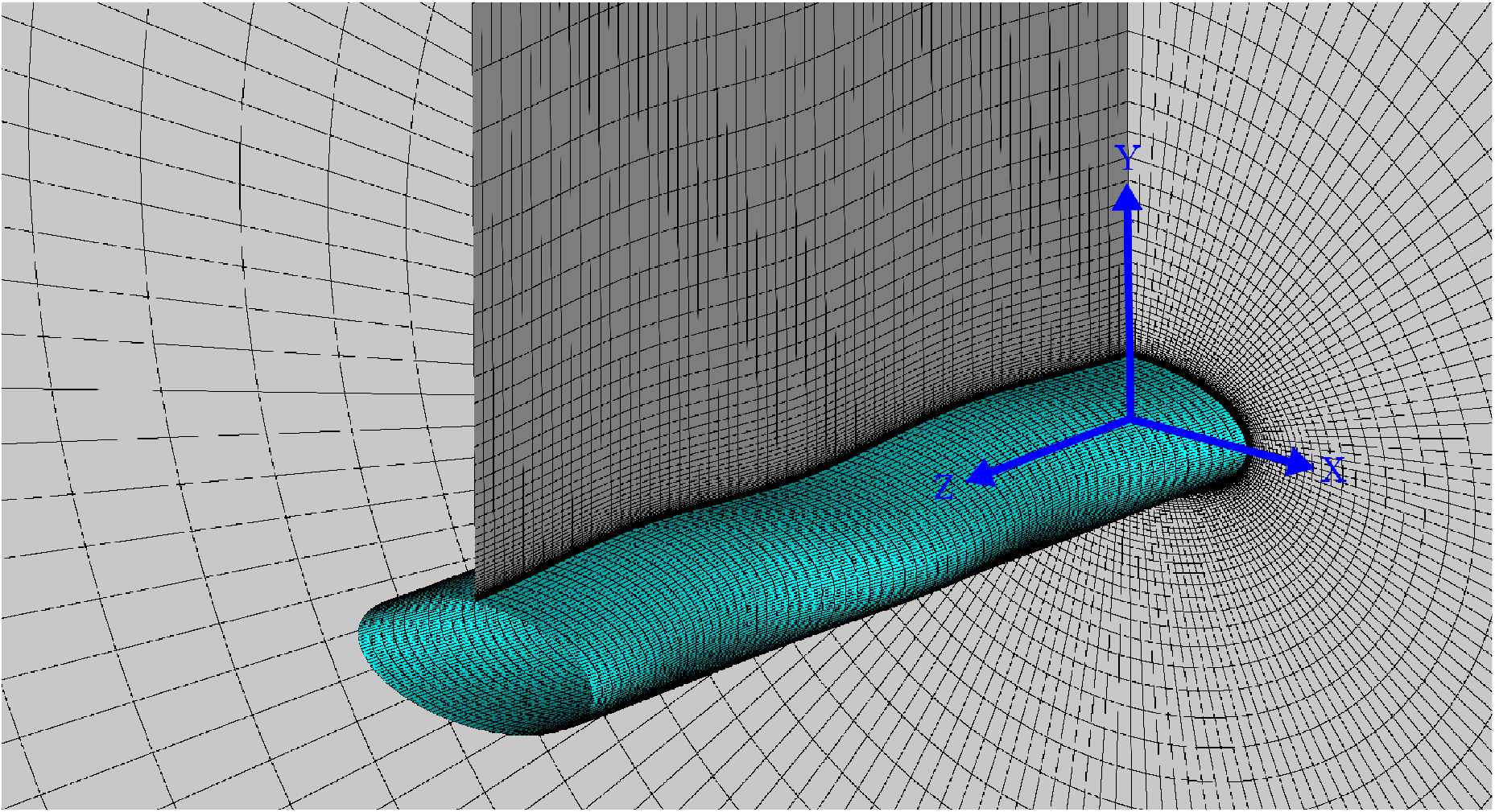
LES mesh for baseline model. Figure shows mesh used for simulation of baseline model using LES at 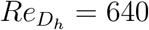 (equivalent to *Re*_*T*_ = 500). Mesh details are listed in Table 4.

Table 3 compares the final mesh selection for the baseline model against a DNS case with a higher resolution mesh, and against other simulations available in the literature. The 16 simulations with varying geometries are all performed at a biologically relevant Reynolds number *Re*_*T*_ = *U*_∞_*T/ν*, of 500, based on the model thickness *T*, which corresponds to a swimming speed of 0.9 m/s [22]. For the baseline model, *Re*_*T*_ = 500 corresponds to 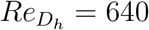, based on the hydraulic diameter *D*_*h*_. To compare directly with the previous literature, two additional simulations are performed at a Reynolds number 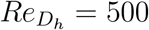. Within this regime, results demonstrate a slight Reynolds number dependence with 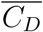 decreasing slightly and *C*_*L,RMS*_ increasing slightly with Reynolds number. The 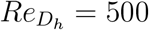 simulations agree very well with the previous literature, and both the resolved DNS with 1.92M cells and the LES at a lower resolution of 0.583M cells have very similar 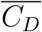 values, within 1% of one another, and both have *C*_*L,RMS*_ values in agreement with previous literature.

**Table 3.** Comparison of mesh variations with previous literature. Various mesh configurations of the baseline model compared with previous literature. *N* is the total size of the computational mesh.

| | Method | $Re_{D_h}$ | $\overline{C_D}$ | $C_{L,RMS}$ | $St$ | $N_{total}$ | $N_z$ | $N_\theta$ | $N_r$ | $\Delta z/T$ | $\Delta r/T_{min}$ |
| --- | --- | --- | --- | --- | --- | --- | --- | --- | --- | --- | --- |
| Current | LES | 500 | 0.749 | 0.0072 | 0.25 | 0.583M | 68 | 130 | 68 | 0.08 | 0.0030 |
| Current | LES | 500 | 0.740 | 0.0081 | 0.21 | 4.85M | 137 | 262 | 137 | 0.04 | 0.0025 |
| Current | DNS | 500 | 0.752 | 0.0092 | 0.22 | 1.92M | 100 | 196 | 100 | 0.07 | 0.0030 |
| Current* | LES | 640 | 0.715 | 0.0141 | 0.20 | 0.718M | 68 | 160 | 68 | 0.10 | 0.0030 |
| Current | DNS | 640 | 0.717 | 0.0105 | 0.22 | 1.92M | 100 | 196 | 100 | 0.07 | 0.0030 |
| Witte et al. [12] | DNS | 500 | 0.769 | 0.0173 | 0.21 | 8M | – | – | – | – | – |
| Hans et al. [14] | LES | 500 | – | 0.011 | 0.2 | – | – | – | – | – | – |
| Kim and Yoon [10] | LES | 500 | 0.741 | 0.0167 | 0.22 | 3.5M | – | – | – | – | – |
| Morrison et al. [11] | DNS | 500 | 0.564 | 0.019 | 0.22 | 55.2M<sup>1</sup> | – | – | – | – | – |

\* representative of final meshing algorithm used in geometry modifications
<sup>1</sup> lattice Boltzmann method

In addition to the quantitative comparison provided in Table 3, the time-averaged streamwise velocity contours are examined in Fig 5, comparing the lower resolution LES to the DNS results. The dominant features of the flow are being captured at the two cross-sections shown. The mean recirculation length along the cross-section, the mean value of the velocities, and the thickness of the recirculation region are consistent between the two simulations. One may note that the mesh chosen for this investigation does not capture far downstream flow structures. However, these structures have little effect on the measured force coefficients. The mesh resolution was carefully chosen such that the root-mean-square lift and the average drag values are comparable to those calculated using a higher resolution mesh. The results of the mesh comparison show average drag values are within 1% of the higher resolution DNS case and root-mean-square lift values differ on the order of 0.004, a smaller deviation than any considered to draw conclusions in this investigation.

**Fig 5.**
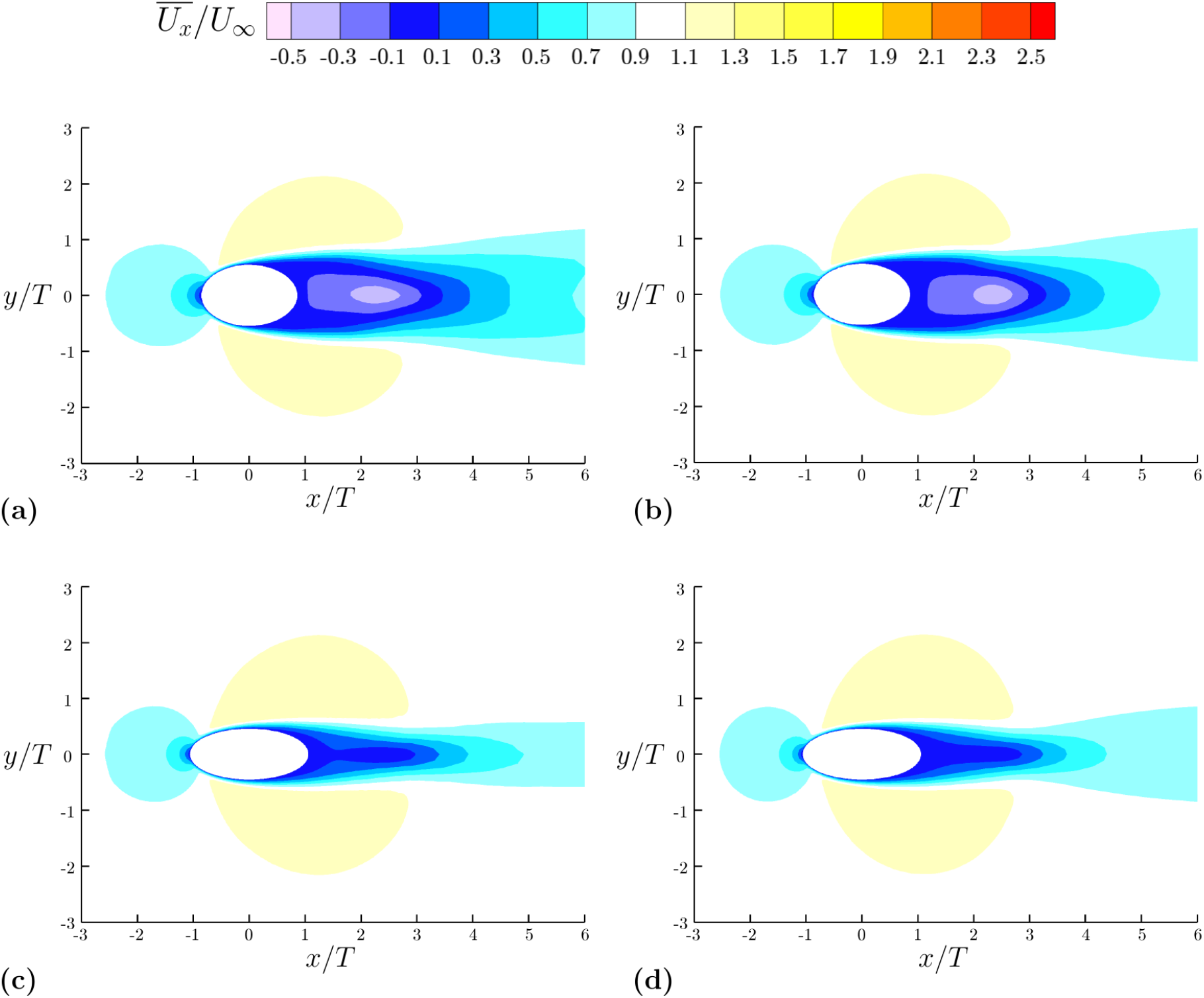
Contours of time-averaged streamwise velocity. Comparison of LES model with fully resolved DNS at 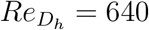 (equivalent to *Re*_*T*_ = 500). Contours of time-averaged streamwise velocity are shown at peak and trough cross-sections of the baseline model. **(a)** LES peak cross-section. **(b)** DNS peak cross-section. **(c)** LES trough cross-section. **(d)** DNS trough cross-section.

The LES with 0.718M points is chosen as the best baseline configuration. The meshes for the remaining 16 simulations follow similar trends as that for the baseline model, however modifications are made to account for the two different aspect ratios (circular and elliptical), and for the two different wavelengths that modify the total spanwise domain of the simulation (long and short). The mesh characteristics for each of these categories are given in Table 4. There are three cases (CS3, ES2, ES4) in which additional mesh points are added to account for highly complex geometries.

**Table 4.** Mesh variation by geometry class. Additional mesh points are added to specific models (CS3, ES2, ES4) within these categories due to increased geometrical complexity.

| Class | $L_z = 2\lambda$ | $\gamma$ | $\lambda$ | $N_{total}$ | $N_z$ | $N_\theta$ | $N_r$ | $\Delta z/T$ | $\Delta r/T_{min}$ |
| --- | --- | --- | --- | --- | --- | --- | --- | --- | --- |
| Circular Short (CS) | 3.6 | 1.0 | 1.8 | 0.718M | 68 | 160 | 68 | 0.054 | 0.0030 |
| Elliptical Short (ES) | 3.6 | 3.0 | 1.8 | 1.01M | 68 | 224 | 68 | 0.054 | 0.0030 |
| Circular Long (CL) | 10.0 | 1.0 | 5.0 | 1.08M | 102 | 160 | 68 | 0.100 | 0.0030 |
| Elliptical Long (EL) | 10.0 | 3.0 | 5.0 | 1.52M | 102 | 224 | 68 | 0.100 | 0.0030 |

## Results and Flow Physics

### Identifying Dominant Geometric Parameters

The results of all 16 geometry modifications in terms of the value of each of the response variables are tabulated in Table 5. Each row also contains the combination of low (−1) and high (1) geometry modifications for the specific configuration (see Table 2). The average effects of the geometric parameters on each response variable are calculated using Eq 5 and Eq 6 and displayed in the Pareto charts in Fig 6.

**Table 5.** Measured response variables by model and geometric variation. See Table 2 for specific geometric values of each variation. Models are grouped by class: Circular Short (CS), Circular Long (CL), Elliptical Short (ES), Elliptical Long (EL).

| Model | $\overline{C_D}$ | $C_{L,RMS}$ | $St$ | $\gamma$ | $\lambda$ | $A_C$ | $A_T$ | $\epsilon$ | $\phi$ |
| --- | --- | --- | --- | --- | --- | --- | --- | --- | --- |
| Baseline | 0.715 | 0.014 | 0.20 | 0 | 0 | 0 | 0 | 0 | 0 |
| CS1 | 1.17 | 0.311 | 0.21 | -1 | -1 | -1 | -1 | -1 | -1 |
| CS2 | 1.10 | 0.100 | 0.21 | -1 | -1 | -1 | 1 | -1 | 1 |
| CS3 | 1.04 | 0.036 | 0.23 | -1 | -1 | 1 | -1 | 1 | 1 |
| CS4 | 1.04 | 0.038 | 0.17 | -1 | -1 | 1 | 1 | 1 | -1 |
| CL1 | 1.05 | 0.075 | 0.22 | -1 | 1 | -1 | -1 | 1 | -1 |
| CL2 | 1.10 | 0.057 | 0.18 | -1 | 1 | -1 | 1 | 1 | 1 |
| CL3 | 1.05 | 0.052 | 0.20 | -1 | 1 | 1 | -1 | -1 | 1 |
| CL4 | 1.02 | 0.037 | 0.12 | -1 | 1 | 1 | 1 | -1 | -1 |
| ES1 | 0.657 | 0.051 | 0.35 | 1 | -1 | -1 | -1 | 1 | 1 |
| ES2 | 0.668 | 0.028 | 0.35 | 1 | -1 | -1 | 1 | 1 | -1 |
| ES3 | 0.655 | 0.044 | 0.33 | 1 | -1 | 1 | -1 | -1 | -1 |
| ES4 | 0.667 | 0.020 | 0.33 | 1 | -1 | 1 | 1 | -1 | 1 |
| EL1 | 0.635 | 0.020 | 0.36 | 1 | 1 | -1 | -1 | -1 | 1 |
| EL2 | 0.633 | 0.008 | 0.22 | 1 | 1 | -1 | 1 | -1 | -1 |
| EL3 | 0.634 | 0.017 | 0.36 | 1 | 1 | 1 | -1 | 1 | -1 |
| EL4 | 0.641 | 0.003 | 0.23 | 1 | 1 | 1 | 1 | 1 | 1 |

**Fig 6.**
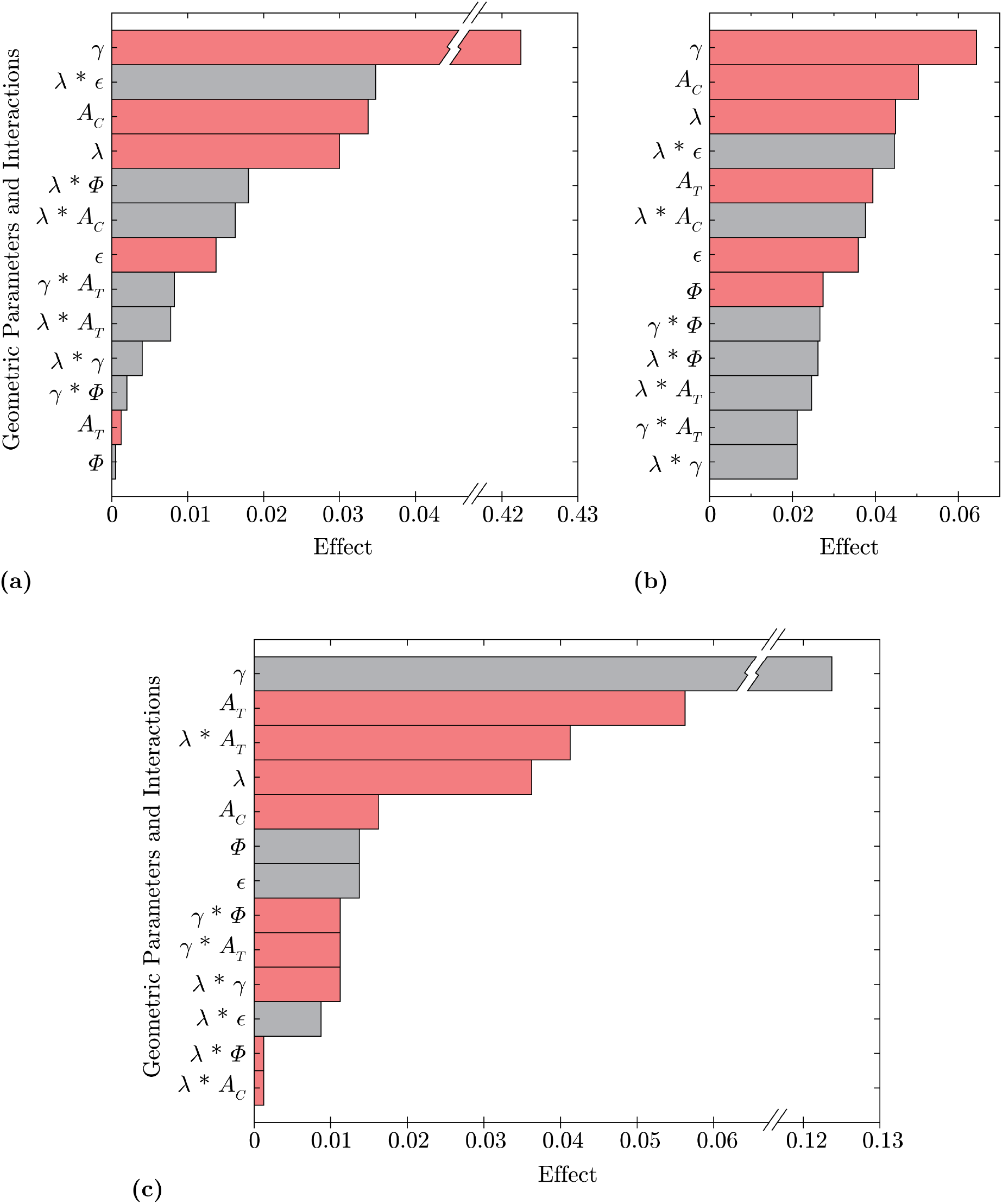
Pareto charts of effects. Three Pareto charts summarize the effects of modifications to geometric parameters on each of the response variables. Gray shaded regions indicate a positive correlation and the red shaded regions indicate a negative correlation. *γ*, *A*_*C*_, *A*_*T*_, and *λ* are seen to have the largest effects. **(a)** 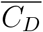 Response **(b)** *C*_*L,RMS*_ Response **(c)** *St* Response.

The Pareto charts rank the effects for each of the three response variables. Most important to the screening test is the overall ranking of effects and not the absolute effect magnitude, however the direction of correlation is indicated by the shading of red (positive correlation) or gray (negative correlation). Interacting parameters have a larger influence together than either does individually, however not every combination of two-parameter interaction is captured since the cumulative effects of multiple two-way interactions are measured simultaneously and each effect cannot be extracted individually, leaving the two-way interactions confounded with one another [26, 27]. However, the test matrix is specifically designed to capture the *λ* and *γ* interactions reported in Fig 6.

Fig 6 shows that the parameters with the largest effect on the response variables are *γ*, *A*_*C*_, *A*_*T*_, and *λ*. The two-way interactions *λ*\**ϵ* and *λ*\**A*_*T*_ are also shown to be significant. In terms of single parameters, the undulation offset, *ϵ*, and the undulation symmetry, *ϕ*, are the least significant. Comparing the response variables to one another, there are more parameters that significantly affect *C*_*L,RMS*_ in contrast to the fewer influential parameters for 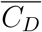. As *C*_*L,RMS*_ is a measure of an oscillating component, it may be more sensitive to changes in geometric parameters than a mean value such as *C*_*D*_. However, the most important factors affecting *C*_*L,RMS*_ are the same as those influencing 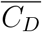, that is *γ*, *A*_*C*_, and *λ*, although, the relative effect of *γ* is smaller with respect to *C*_*L,RMS*_ than in the 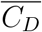 response.

Although *γ* is the highest ranking parameter on all of the Pareto charts, its strong effect on the response parameters is not surprising as the influence of aspect ratio is well known in aerodynamics. The high and low values of *γ* distinguish between the two classes of geometries tested, those with a baseline circular aspect ratio and those with a streamlined elliptical aspect ratio. The other undulation parameters can be applied to each of these regimes, and their impacts are discussed below by identifying simulations with high and low parameter values within a specific aspect ratio class.

### Effect of *A*_*T*_

To analyze the effect of a change in thickness amplitude, *A*_*T*_, two similar geometries with varying *A*_*T*_ attributes are compared in Fig 7, with EL1 (low *A*_*T*_) in Fig 7a and EL2 (high *A*_*T*_) in Fig 7b. The two geometries have a similar *C*_*D*_, however, the high undulation amplitude of EL2 reduces the *C*_*L,RMS*_ by 60%.

**Fig 7.**
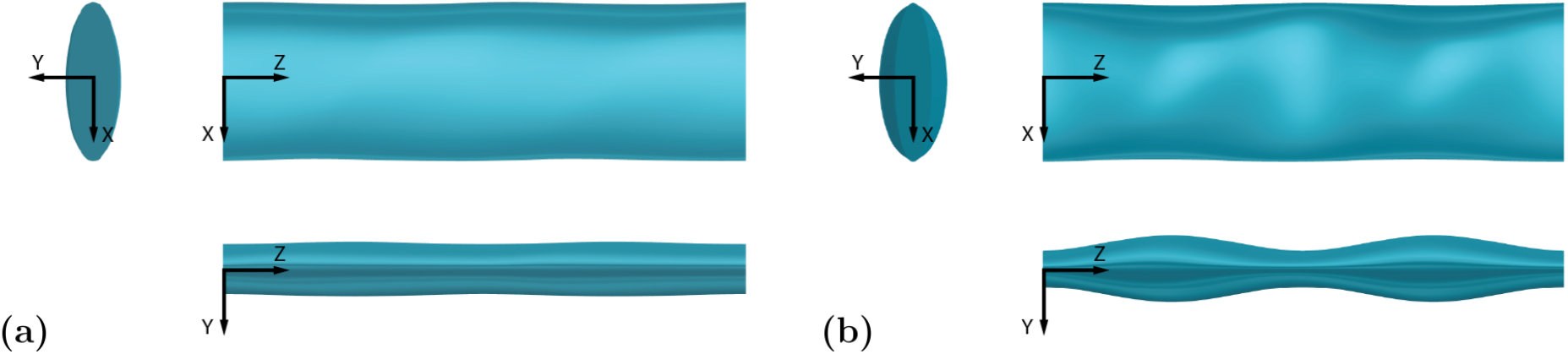
Geometry of models with low and high *A*_*T*_ values. Directly comparing a low *A*_*T*_ and high *A*_*T*_ model with three views: top view in the *x-z* plane, front view in the *y-z* plane, and side view in the *x-y* plane. Flow is along the positive *x*-axis. **(a)** EL1 (low *A*_*T*_). **(b)** EL2 (high *A*_*T*_).

The time-averaged streamwise velocity contours in Fig 8a show considerable differences between the two geometries. The recirculation length across the peak and trough locations are nearly constant for EL1, whereas they vary dramatically across the span for EL2. EL2 has substantial flow reversal at peak thickness in contrast to the trough location where the recirculation length is dramatically reduced due to a locally reduced frontal area. The streamlines in Fig 8b also show little to no recirculation immediately downstream of the trough cross-section of EL2 and high spanwise velocities downstream of the peaks. The lack of a uniform velocity profile between peak and trough regions promotes spanwise momentum transport and results in the vortex break up shown in Fig 8c, which displays isosurfaces of the *Q*-criterion colored by magnitude of spanwise vorticity. Positive vorticity values (counter-clockwise rotation) are displayed in red, while negative values (clockwise rotation) are displayed in blue.

**Fig 8.**
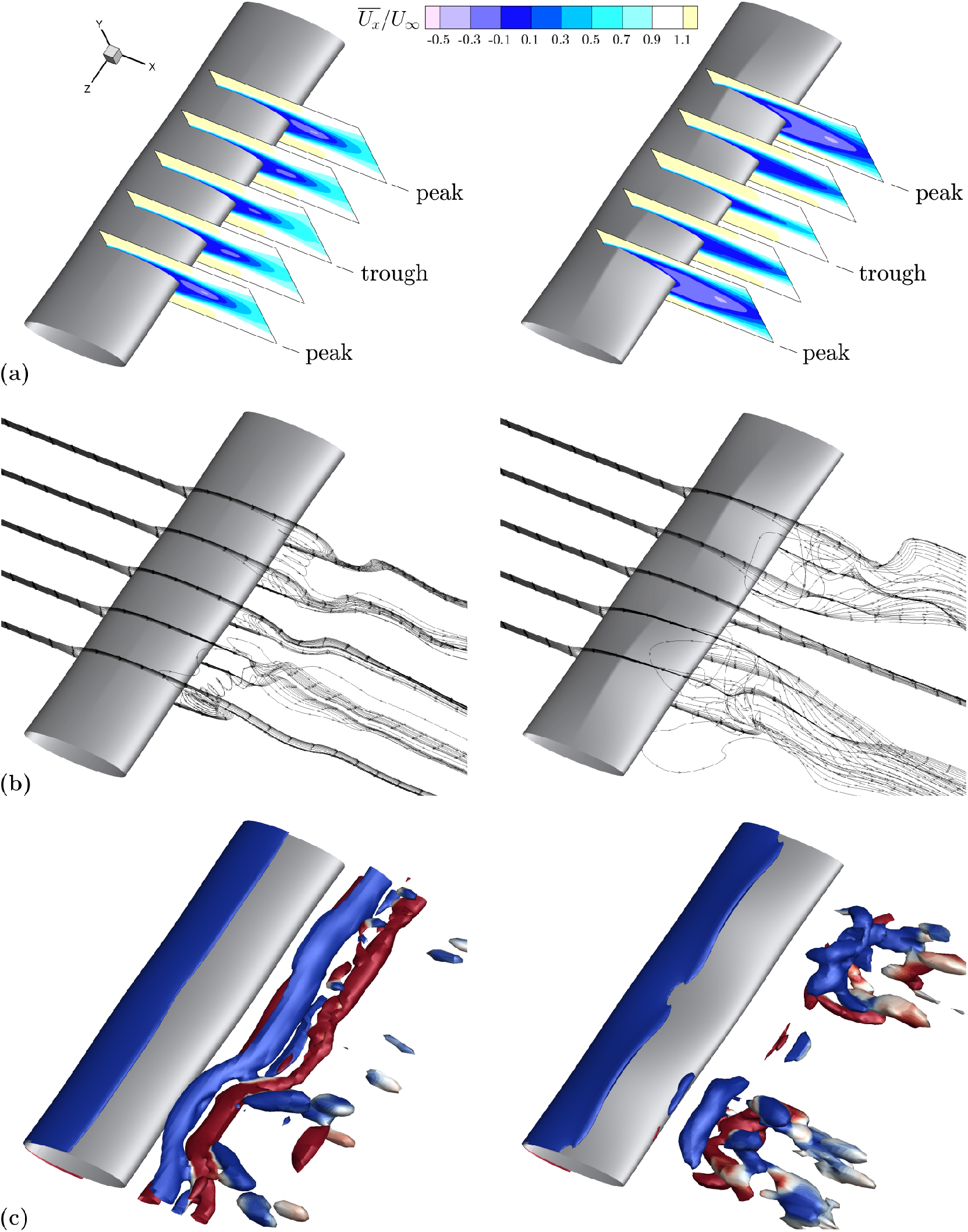
Flow comparison between low and high *A*_*T*_ geometries. EL1 (low *A*_*T*_) in the left column, EL2 (high *A*_*T*_) in the right column. **(a)** The comparison of time-averaged streamwise velocity contours for low *A*_*T*_ (left) and high *A*_*T*_ (right) models shows considerable variation between recirculation length at peak and trough cross sections. Contours are shown at equally spaced spanwise locations where peak and trough correspond to *A*_*T*_ undulations. **(b)** Velocity streamlines at peak and trough spanwise locations denoted in 8a demonstrate more spanwise transport for the high *A*_*T*_ model. **(c)** Isosurfaces of nondimensional *Q*-criterion value 0.8, colored by positive (red) and negative (blue) *z*-vorticity. Isosurfaces display long, coherent structures at low *A*_*T*_ and breakup at high *A*_*T*_.

Fig 9 displays the *C*_*L*_ frequency spectra for EL1 and EL2 compared with a smooth elliptical cylinder of the same aspect ratio. The EL1 (low *A*_*T*_) and cylinder spectra show a similarity in shape and peak value around *f*\* = 0.35. EL1 also has a secondary peak around *f*\* = 0.22 not present for the smooth cylinder. At high *A*_*T*_, the *f*\* = 0.35 peak is attenuated and the *f*\* = 0.22 peak becomes dominant. The attenuation in the *f*\* = 0.35 peak leads to a lower overall signal magnitude evident in the strong reduction of *C*_*L,RMS*_ for high *A*_*T*_. The difference in dominant shedding frequencies is likely due to the change in development, growth, and shedding of the differently sized and shaped spanwise structures as indicated in the isosurfaces in Fig 8c.

**Fig 9.**
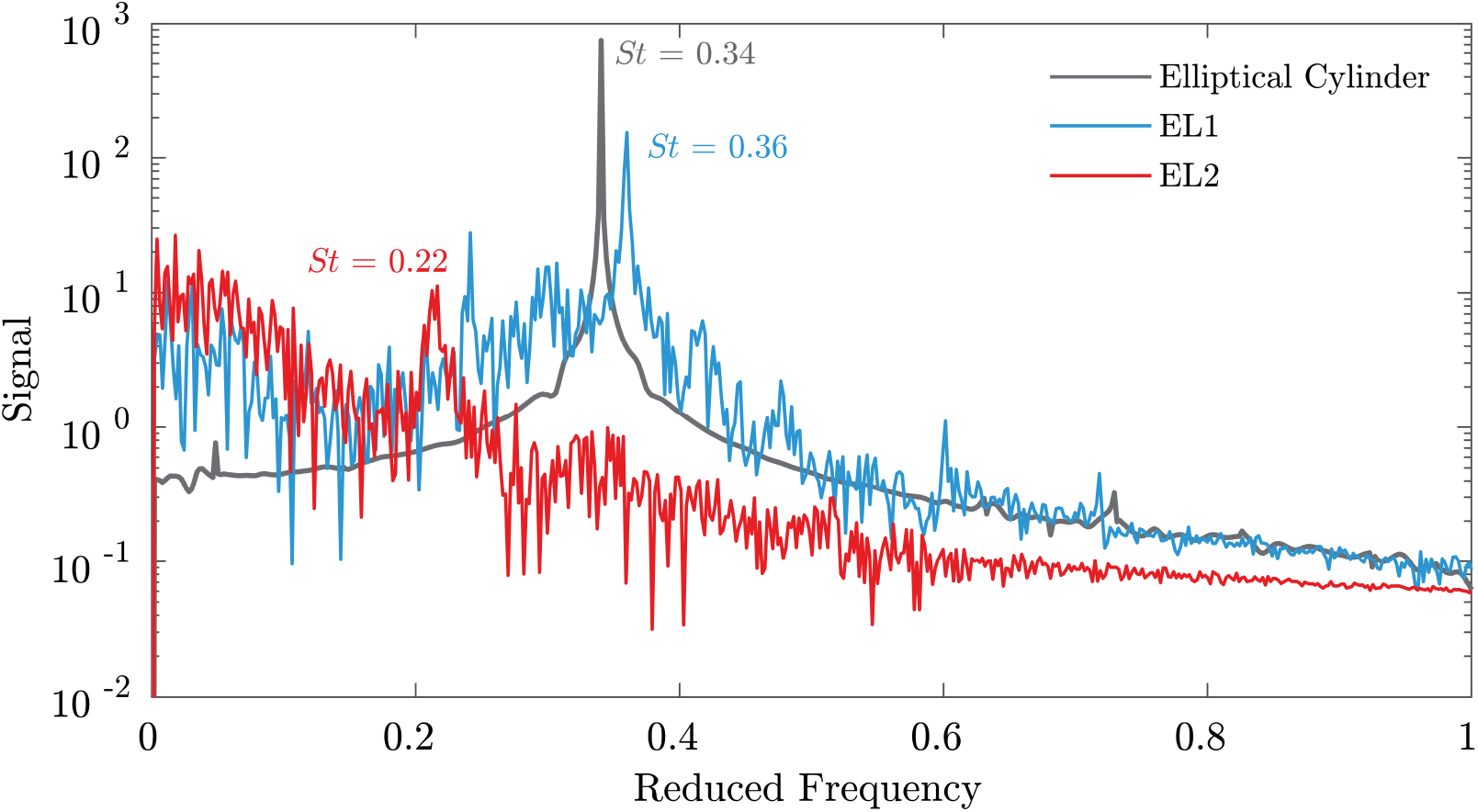
*C*_*L*_ spectra for low and high *A*_*T*_ geometries. The *C*_*L*_ spectra for EL1 (low *A*_*T*_) and EL2 (high *A*_*T*_) geometries shown in Fig 7 are directly compared with a smooth elliptical cylinder. The EL1 frequency spectrum shows a spectra more similar to the cylindrical case while the EL2 response is smaller in magnitude with a peak at lower frequency.

### Effect of *A*_*C*_

The effect of the chord length amplitude, *A*_*C*_, is shown by comparing the flow over models CL2 (low *A*_*C*_) and CL4 (high *A*_*C*_) in Fig 10. These two geometries both have high *A*_*T*_, high *λ*, and a circular aspect ratio. As *A*_*C*_ is increased, the response parameters show a 7% reduction in 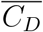 and a 35% decrease in *C*_*L,RMS*_.

**Fig 10.**
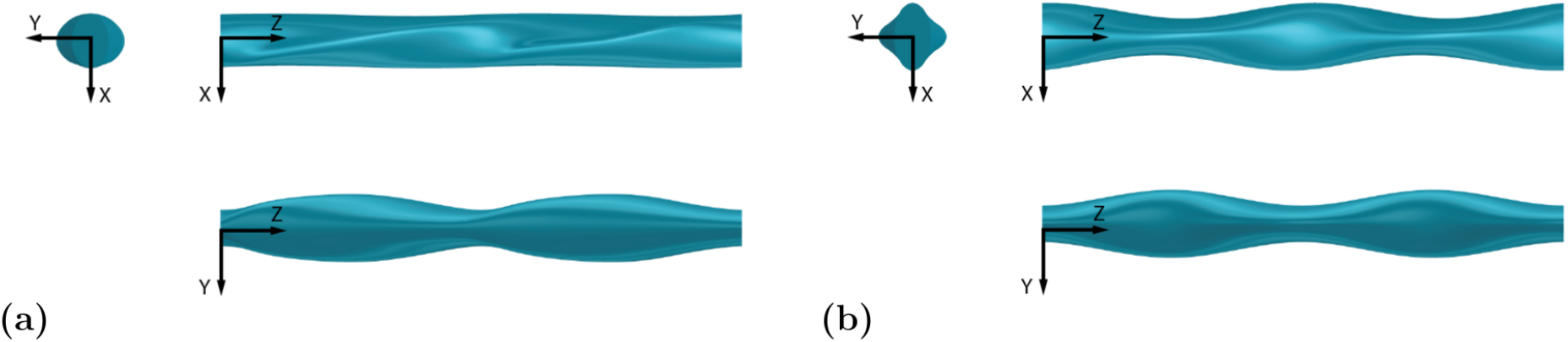
Geometry of models with low and high *A*_*C*_ values. Directly comparing a low *A*_*C*_ and high *A*_*C*_ model with three views in the same configuration as Fig 7. **(a)** CL2 (low *A*_*C*_). **(b)** CL4 (high *A*_*C*_).

Contour plots of the mean streamwise velocity and associated streamlines are shown in Fig 11a and Fig 11b. Along the span, both models have a similar pattern of increasing and decreasing recirculation lengths, primarily the result of their similarly high *A*_*T*_ parameter as previously discussed. However, differences are seen in the *Q*-criterion isosurfaces in Fig 11c where the high *A*_*C*_ geometry has no coherent vortex structure in the center of the span where the chord length is the largest. The local, more elliptical, cross-section mitigates flow separation in the center of the span, thus reducing drag and lift fluctuations as *A*_*C*_ is increased. The effect of increasing *A*_*C*_ has a larger impact on circular aspect ratio models. For these geometries, the change in the chord length due to the high *A*_*C*_ value in Table 2 represents a larger percent of the overall chord length than for elliptical aspect ratio models.

**Fig 11.**
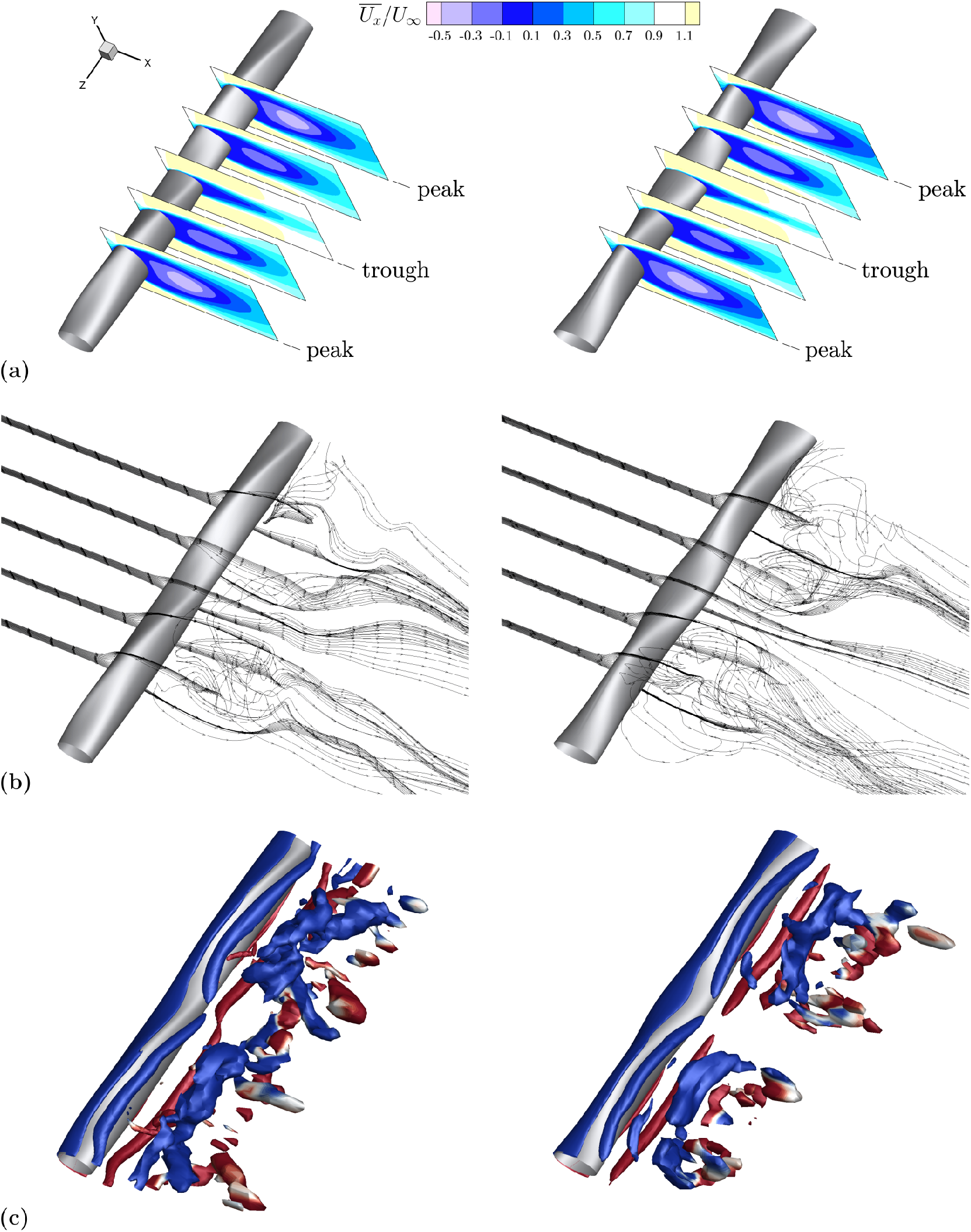
Flow comparison between low and high *A*_*C*_ geometries. CL2 (low *A*_*C*_) in the left column, CL4 (high *A*_*C*_) in the right column. **(a)** The comparison of time-averaged streamwise velocity contours for low *A*_*C*_ (left) and high *A*_*C*_ (right) models shows a relative similarity between the two models. Contours are shown at equally spaced spanwise locations where peak and trough correspond to *A*_*T*_ undulations. **(b)** Velocity streamlines at peak and trough spanwise locations denoted in 11a demonstrate the similarity in the streamwise flow component and a modest increase in spanwise flow for the high *A*_*C*_ model. **(c)** Isosurfaces of nondimensional *Q*-criterion value 1.6, colored by positive (red) and negative (blue) z-vorticity. Isosurfaces display a noticeable break towards the center of the high *A*_*C*_ model.

The *C*_*L*_ spectra for CL2 and CL4 shown in Fig 12 show that the intermittent spanwise features are correlated with a shift to lower frequencies and lower amplitudes. The smooth circular cylinder has the strongest and highest frequency peak, that gradually shifts to *St* = 0.18 for low *A*_*C*_ and to *St* = 0.12 for high *A*_*C*_. Moreover, the high *A*_*C*_ model has reduced amplitudes at *f*\* > 0.6 and an overall lower magnitude at *St* = 0.12, consistent with the reduction in *C*_*L,RMS*_.

**Fig 12.**
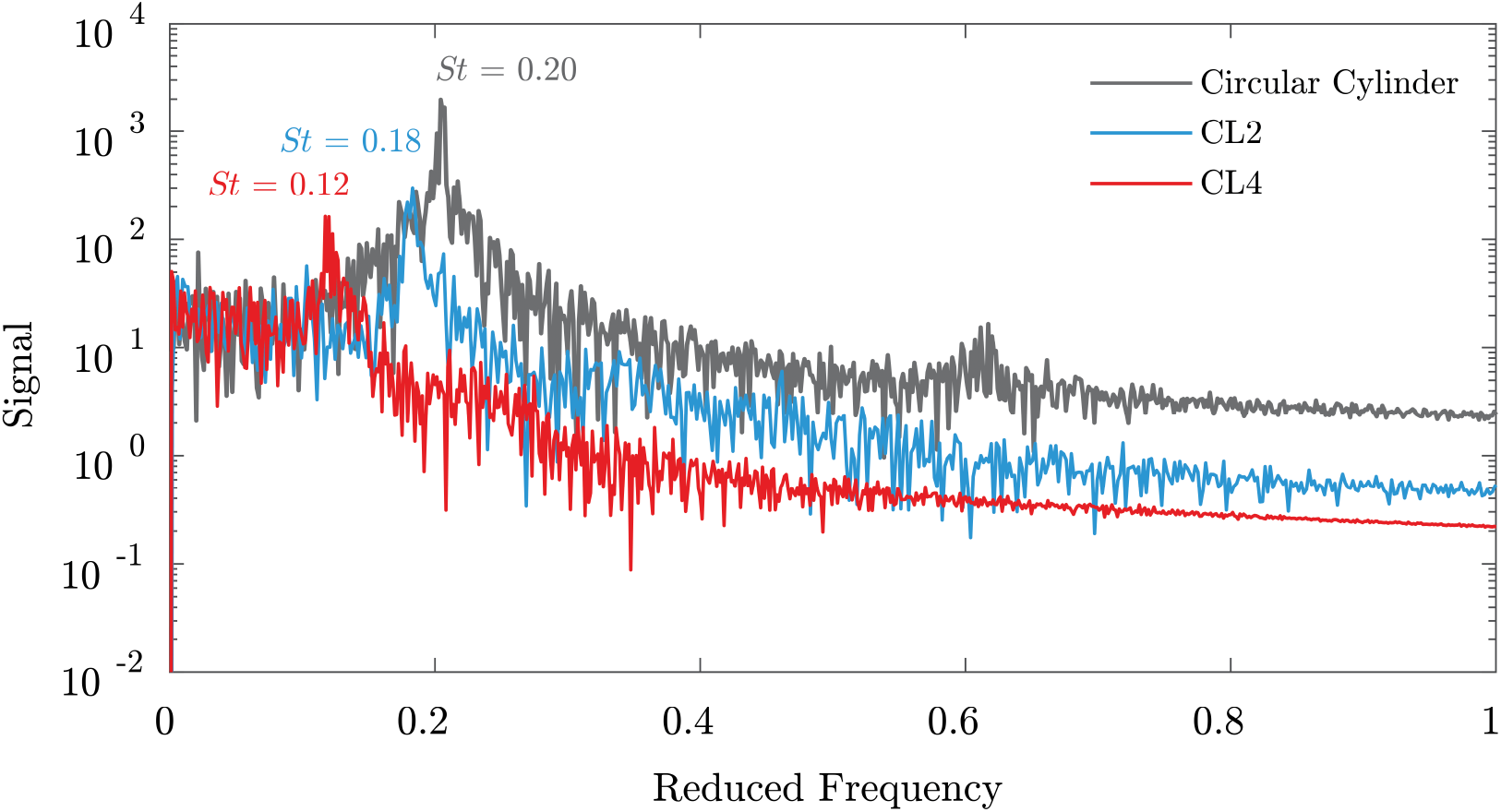
*C*_*L*_ spectra for low and high *A*_*C*_ geometries. The *C*_*L*_ spectra for the CL2 (low *A*_*C*_) and CL4 (high *A*_*C*_) geometries shown in Fig 10 are directly compared with a smooth circular cylinder. A gradual decrease in *St* occurs as *A*_*C*_ increases.

### Effect of *λ*

*Q*-criterion isosurfaces for cases with low and high wavelength are presented in Fig 13 for both circular and elliptical aspect ratio models and directly compared to smooth cylinders. The vortex structures shed from the low wavelength models take on the appearance of long spanwise coherent vortices, qualitatively similar to the smooth structures despite the large value of *A*_*T*_. In contrast, the high *λ* case demonstrates considerable spanwise break up of the downstream vortex structures for both cross-sections.

**Fig 13.**
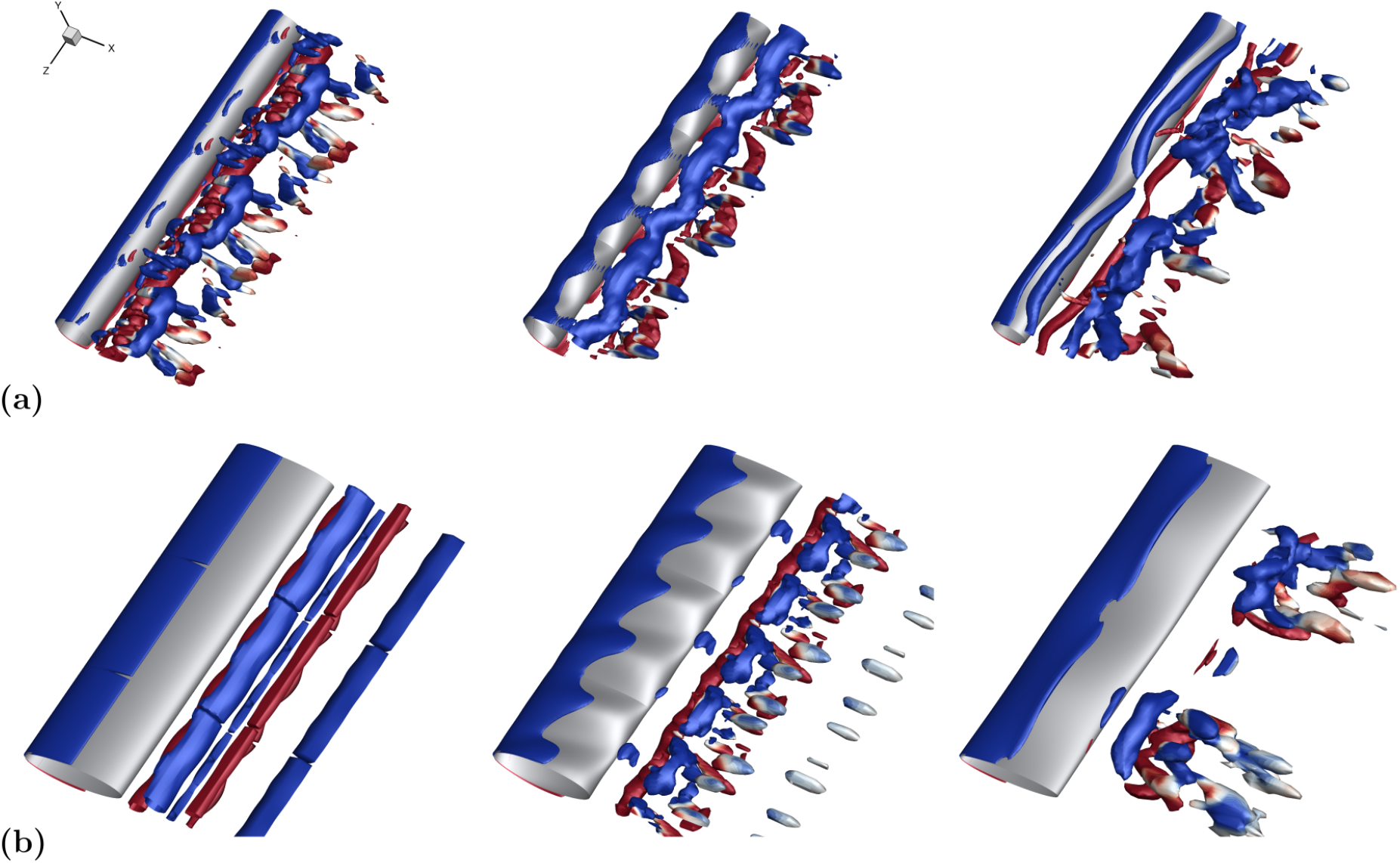
Flow structure comparison among various *λ* geometries. Isosurfaces of *Q*-criterion colored by positive (red) and negative (blue) spanwise vorticity show a larger wavelength allows for break up of flow structures while flow structures from low *λ* models continue to resemble those of their smooth cylindrical counterparts. **(a)** Circular geometries (low *γ*); nondimensional *Q*-criterion value 1.6. From left to right: no undulations, CS2 (low *λ*), CL2 (high *λ*). **(b)** Elliptical geometries (high *γ*); nondimensional *Q*-criterion value 0.8. From left to right: no undulations, ES2 (low *λ*), EL2 (high *λ*).

In the *C*_*L*_ spectra shown in Fig 14, the low *λ* cases have *St* peaks close to the frequency of their smooth cylinder counterparts, whereas the high *λ* cases have an appreciable decrease in *St*. Increasing the wavelength also increases the magnitude of the shift and decreases the amplitude of the peak, paralleling the reduction in *C*_*L,RMS*_. The effect of the interaction *λ*\**A*_*T*_, noted in the *St* Pareto chart (Fig 6c), is seen here. For high values of both *λ* and *A*_*T*_, the reduction in *St* is greatly increased.

**Fig 14.**
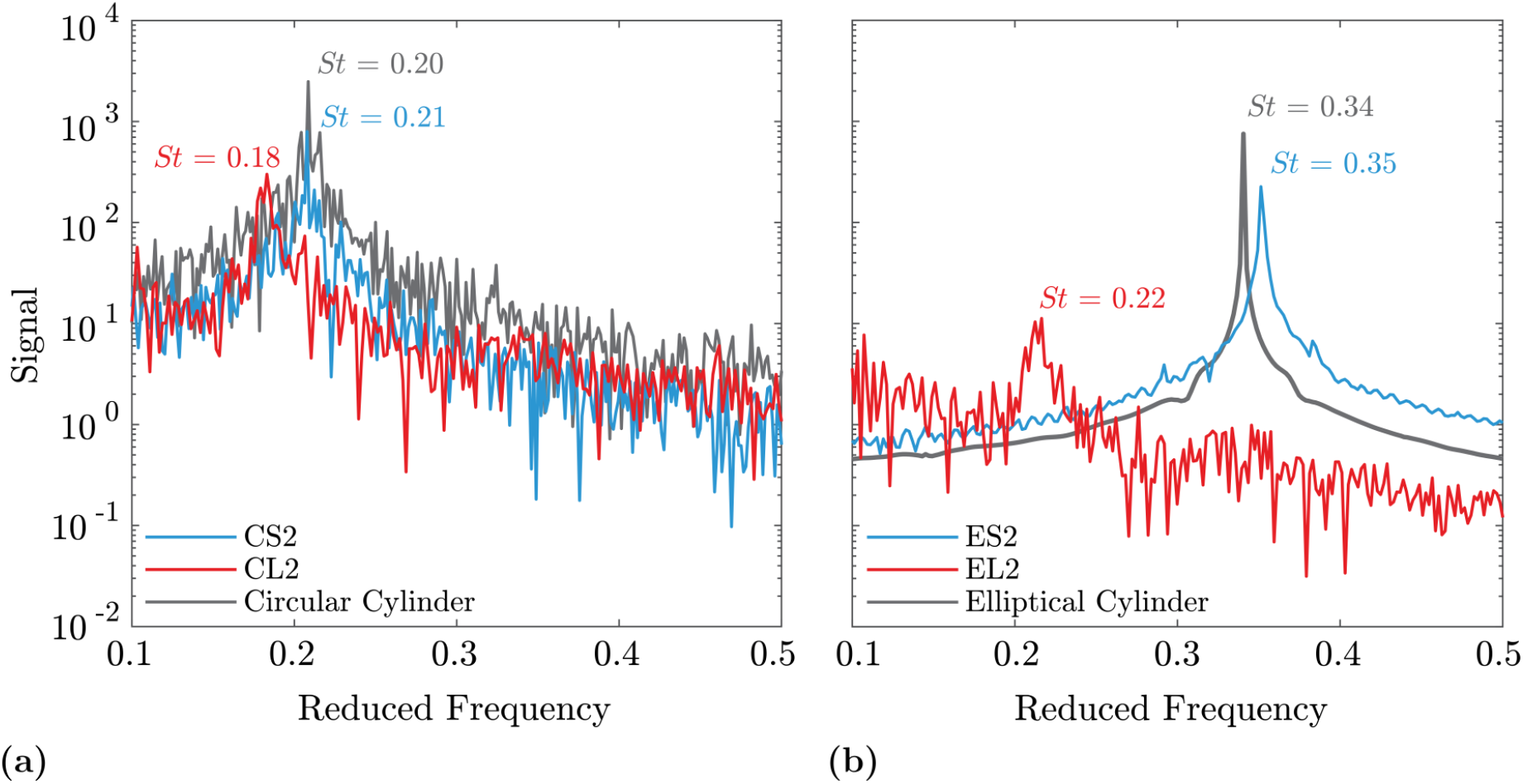
Frequency spectra for low and high *λ* geometries. The frequency spectra for low and high values of *λ* and *γ* are compared with the smooth cylinder cases showing *St* similarity between low *λ* and smooth models. **(a)** Circular aspect ratio. **(b)** Elliptical aspect ratio.

### Effect of Undulation Offset (*ϵ*) and Symmetry (*ϕ*)

The effects of *ϵ* and *ϕ* are less prominent, and their impact is primarily overshadowed by the previously discussed geometric parameters. Nevertheless, the appearance of a large value of *ϵ* can incrementally decrease 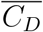 and *C*_*L,RMS*_. As streamwise flow travels along the chord length, the offset of the undulations increases the breakup of the spanwise structures, especially for geometries with a high *A*_*C*_ value, although the interaction *A*_*C*_\**ϵ* is not specifically computed. The impact of other geometric parameters is thereby augmented by the high value of *ϵ*. The interaction *λ*\**ϵ* is also shown to be important to both 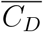 and *C*_*L,RMS*_. The complexity of this relationship is not fully understood and mostly likely results from the variety in flow patterns induced by changes in wavelength.

Based on the current set of simulations, little quantitative data exists to support a correlation between *ϕ* and any of the calculated response variables. However, the parameter does impact the flow field and is most apparent for high *A*_*C*_ models. A high *ϕ* preferentially skews the spanwise velocity towards one direction as shown by model EL4 in Fig 15. The model shown has a high value of *A*_*T*_, *A*_*C*_, *ϵ*, and *ϕ*. As discussed previously, a high *A*_*T*_ value results in considerable momentum transport in the *z*-direction. The further addition of high *ϵ* and *ϕ* values results in this spanwise motion being directed nearly entirely in the negative *z*-direction. The asymmetry can be seen in the streamlines shown in Fig 15b as well as the time-averaged *z*-velocity contour in Fig 15d as the shed vortical structures tend towards the negative *z*-direction.

**Fig 15.**
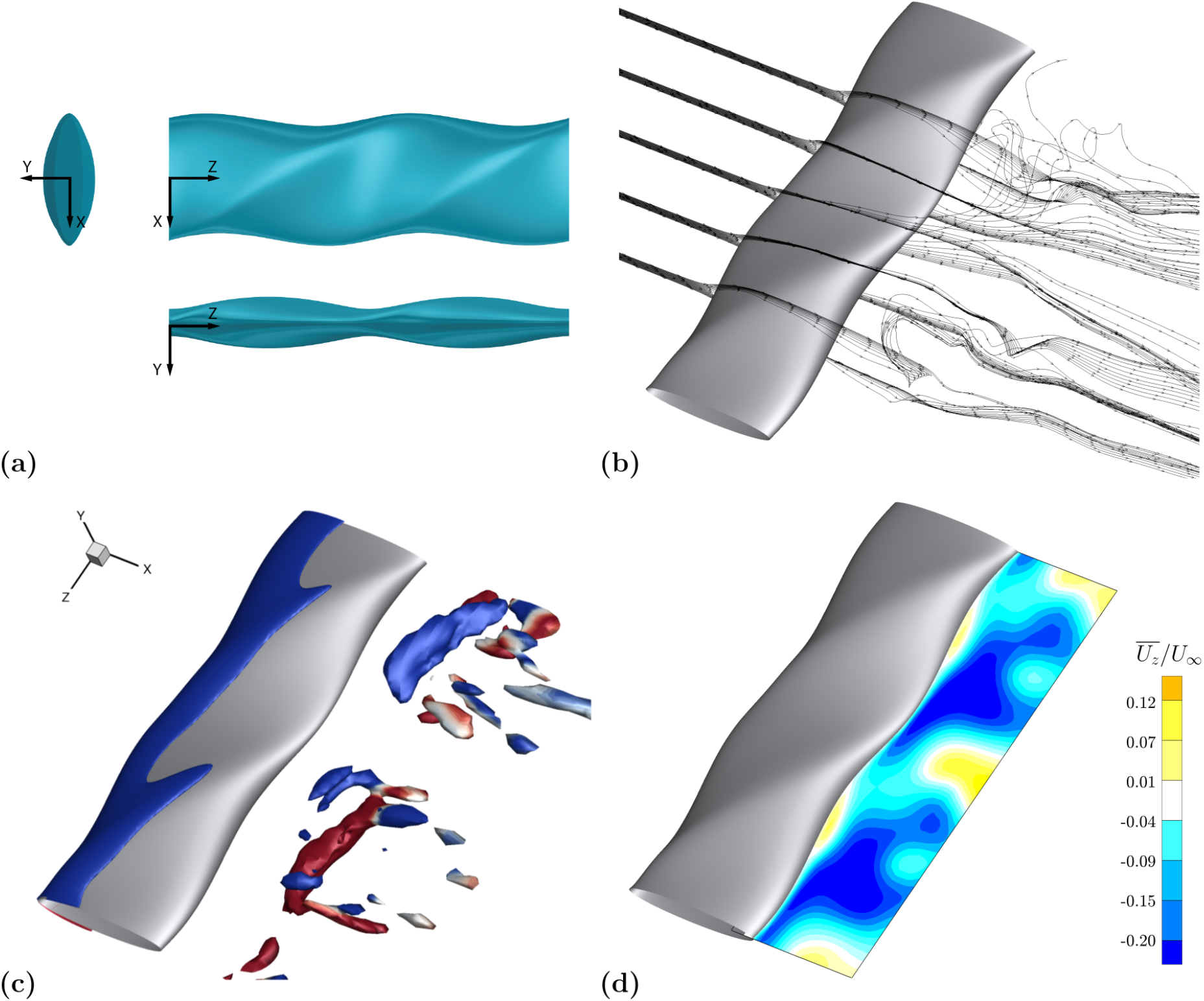
Flow visualization for high *ϕ* and *ϵ* geometry. Flow visualization reveals disjointed vortex structures and a preferential direction for the spanwise velocity. **(a)** EL4 (high *ϵ* and high *ϕ*): top view in the *x-z* plane, front view in the *y-z* plane, and side view in the *x-y* plane. **(b)** Velocity streamlines at equally spaced spanwise locations as described in Fig 11. **(c)** Isosurfaces of nondimensional *Q*-criterion value 0.8, colored by positive (red) and negative (blue) *z*-vorticity. **(d)** Mean spanwise velocity in *x-z* plane.

## Discussion

The combination of thickness undulations and chord length undulations is unique to the seal whisker geometry. Consistent with the findings of Hans et al. [14], the magnitudes of both of these undulations, *A*_*T*_ and *A*_*C*_, are shown to reduce *C*_*L,RMS*_. Modification to *A*_*T*_ also influences the frequency response, shifting the *St* to lower values for high *A*_*T*_ geometries. The current data indicate that an increase in *A*_*T*_ alone is capable of shifting *St* to lower frequencies and lower amplitudes by breaking up the strong coherent vortex structures characteristic of smooth circular cylinders. This vortex break up is explained by an increase in spanwise flow velocity at peak thickness locations, resulting in a subsequent decrease in oscillatory transverse flow and a reduction in *C*_*L,RMS*_. Increasing *A*_*C*_ also reduces oscillations in lift, however through a different mechanism. A high *A*_*C*_ value creates cross-sections of varying chord lengths along the span, and thus the flow remains attached at locations of maximum chord length, considerably impacting the local drag for that portion of the span. Although the streamlines still separate dramatically on spanwise portions with more circular cross-sections, the regions of attached flow drive down both *C*_*L,RMS*_ as well as 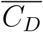.

Indeed, the current results suggest that *A*_*C*_ more strongly influences the drag response compared to *A*_*T*_. This result differs from the observations of Hans et al., who report that thickness undulations have a higher impact on drag than chord length undulations [14]. The discrepancy is likely due to different geometric testing methodologies. Hans et al. chose to eliminate one direction of undulation at a time, whereas the current method draws conclusions from a set including a low and a high value of each amplitude. Both methodologies only examine a small portion of the parameter space in terms of amplitudes, and there are certainly regimes and subtleties that are not fully captured and are worthy of future investigation.

The periodicity of the seal whisker geometry is characterized by the wavelength, *λ*, that governs the frequency of both undulations. The data demonstrates the strong relationship between the flow physics and the wavelength, *λ*, as it has a significant effect on all the response values investigated. The wavelength has been shown to have specific regimes of different flow responses in previous investigations of simpler wavy cylinders (single undulation in all directions) [21]. The different behavior between the low and high wavelength models of the simulation matrix indicate that these regimes may persist in the more complex undulations of the seal whisker model. The spanwise variation of wavy cylinders has been shown to suppress the formation of larger-scale spanwise vorticity [20]. Similar suppression is seen here for large wavelength models. Whereas the low *λ* models maintain coherent spanwise vortex structures as they are shed from the surface, the high *λ* models demonstrate more complex vortex patterns and a substantially different frequency response for the unsteady lift force.

Recent work completed by Liu et al. on seal whisker models also indicates a relationship between wavelength and varying flow regimes [15]. The results presented here, however, include effects of other geometric parameters and suggest a threshold wavelength below which other undulation parameters have limited effect on the flow response. Still, due to the small number of wavelengths tested, the full flow regime map resulting from wavelength changes in the seal whisker topography remains to be determined.

The final two geometric parameters, *ϵ* and *ϕ*, govern the symmetry of the seal whisker geometry. The effect of these geometric parameters are closely related to the inclination angles of the two ellipses, *α* and *β*, introduced by the original seal whisker model (Fig 2). These angles have shown high variability in seal whisker sample measurements and thus have been eliminated from various computational models [12, 15] as inconsequential. Hans et al. show the angles *α* and *β* to have relatively little effect on the oscillatory response. However, when *α* and *β* are decreased, the undulation amplitudes also increase, as the two parameters are coupled. Thus, the novelty of the new geometric definitions introduced in this paper is that the geometric asymmetry parameters *ϵ* and *ϕ* (shown in Fig 3) can both be modified independently from one another and from the two amplitudes. Thus, this investigation can more accurately isolate the dependence of these parameters.

The data indicates that *ϵ* is more important than *ϕ* for all the response parameters, or the undulation offset between leading and trailing edge has a larger impact on the flow response than the symmetry of the undulations. Especially for models with high values of *A*_*C*_, a high value of *ϵ* can incrementally decrease both 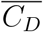 and *C*_*L,RMS*_. Interactions between *λ* and *ϵ* also contribute to the force response and may be pursued in more detailed studies. The parameter *ϕ* has relatively little effect on all three response variables.

## Conclusion

The complex morphology of the harbor seal whisker is modeled with seven independent geometry parameters whose effects on response variables 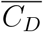, *C*_*L,RMS*_, and *St* are systematically tested. A simulation matrix of 16 seal whisker models, comprised of unique nondimensional geometric parameter combinations, is created using a two-level fractional factorial design of experiments. The evaluation of 16 models is the largest study of geometry modifications to be completed on the seal whisker geometry to the authors’ knowledge. By applying large changes in the independent geometric parameters, the parameters that have a significant impact on 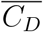, *C*_*L,RMS*_, and *St* are identified as *γ*, *A*_*C*_, *A*_*T*_, and *λ*. The flow physics associated with low and high values of each of these parameters are discussed.

The effects of all the geometric parameters, summarized in Fig 6, are designed to be a preliminary screening tool for more in-depth investigations. The initial flow analyses performed here does indicate specific trends supported by the flow physics, and isolate the dominant geometric features that most influence those trends. These findings are instrumental for bio-inspired design of a variety of structures, small and large, in which drag reduction, vibration suppression, and/or frequency tuning is required. Further research may investigate the effect of other geometric interactions and will likely focus on the optimal range of each of the most important parameters for various implementations.

The impact of understanding the physical response to geometric variations will also be of importance to the biological community. By exaggerating the physical characteristics of the seal whisker beyond that which is observed in nature, the biology community can better identify the importance of morphology and provide context for the natural variation observed within and between species. In future work, a focused study across species may lead to insight into how whisker morphology impacts foraging and prey specialization.

## Acknowledgments

The authors kindly thank Mr. Andrew Guarendi for his assistance in the simulations, and Prof. Kenny Breuer for providing laboratory space. This research was conducted using computational resources and services at the Center for Computation and Visualization, Brown University.

## Distribution Statement A

Approved for public release. Distribution is unlimited.

